# Beyond the Linear Genome: Comprehensive Determination of the Endogenous Circular Elements in *C. elegans* and Human Genomes via an Unbiased Genomic-Biophysical Method

**DOI:** 10.1101/128686

**Authors:** Massa J. Shoura, Idan Gabdank, Loren Hansen, Jason Merker, Jason Gotlib, Stephen D. Levene, Andrew Z. Fire

## Abstract

Investigations aimed at defining the 3-D configuration of eukaryotic chromosomes have consistently encountered an endogenous population of chromosome-derived circular genomic DNA, referred to as extrachromosomal circular DNA (eccDNA). While the production, distribution, and activities of eccDNAs remain understudied, eccDNA formation from specific regions of the linear genome has profound consequences on the regulatory and coding capabilities for these regions. High-throughput sequencing has only recently made extensive genomic mapping of eccDNA sequences possible and had yet to be applied using a rigorous approach that distinguishes ascertainment bias from true enrichment. Here, we define eccDNA distribution, utilizing a set of unbiased topology-dependent approaches for enrichment and characterization. We use parallel biophysical, enzymatic, and informatic approaches to obtain a comprehensive profiling of eccDNA in *C. elegans* and in three human cell types, where eccDNAs were previously uncharacterized. We also provide quantitative analysis of the eccDNA loci at both unique and repetitive regions. Our studies converge on and support a consistent picture in which endogenous genomic DNA circles are present in normal physiological DNA metabolism, and in which the circles come from both coding and noncoding genomic regions. Prominent among the coding regions generating DNA circles are several genes known to produce a diversity of protein isoforms, with mucin proteins and titin as specific examples.

## Introduction

Endogenous DNA circles derived from canonical linear chromosomal loci, known as extrachromosomal circular DNA (eccDNA), were first detected in nuclear fractions of plant cells (wheat and tobacco) in the 1980s by electron microscopy.(Kinoshita et al., 1985) Since then, eccDNAs have been detected in human cell lines(Assum et al., 1989; Cohen et al., 2008; Kinoshita et al., 1985; Kuttler and Mai, 2007; Misra et al., 1989) and cells of various organisms.(Gaubatz, 1990) Accumulating levels of eccDNA have been observed in connection with developmental progression,(Gaubatz and Cutler, 1990; Gaubatz and Flores, 1990) aging,(Gaubatz, 1990; Gaubatz and Flores, 1990; Kinoshita et al., 1985; Sinclair and Guarente, 1997) and genome instability.(Cohen et al.) Differences in eccDNA copy number and/or expression suggest that eccDNAs can contribute to genomic variation and mosaicism in different tissues, expanding the diversity in coding and regulatory capacity of eukaryotic genomes and transcriptomes.

A subset of eccDNA elements are associated with malignancies and drug-resistant tumors in a wide variety of cancers, such ascircularized oncogenes and drug resistance factors (“Double minutes”), which are capable of driving events in oncogenesis.(Albertson et al., 2003; Carroll et al.; Snijders et al., 2003) Beyond the known oncogene and mobile elements, multiple genomically unstable phenotypes are associated with accumulation of eccDNAs,(Cohen et al.; Cohen et al.; Cohen and Lavi; Cohen and Segal; Gaubatz, 1990; Sinclair and Guarente) including an observed rise in eccDNA levels in cells treated with carcinogens (Cohen and Lavi) and in fibroblasts from patients suffering from Fanconi’s anemia.(Cohen et al., 1997) Moreover, deletions of genomic DNA segments in a circular form can occur in programmed processes such as RAG-dependent VDJ recombination at the immunoglobulin and T-Cell receptor loci in vertebrates. It should be noted that somatic deletions are generally investigated only when there is an associated phenotype; therefore, there has been little opportunity to assess the level and scope of DNA deletions and corresponding eccDNAs in healthy cells.

Circular DNA elements are often unrecognized or lost in whole-genome studies that depend on existing tools, even when focused on a specific loci of interest. Thus, eccDNA remains a relatively unexplored component of the eukaryotic genome. (Cesare and Reddel; Cohen and Segal, 2009; Dilley et al.) For instance, despite having tremendous progress in our understanding of the sequence and the 3D structure of eukaryotic linear genomes and the mechanisms of dsDNA breakpoint repair, not much is known about the fate of the deleted/excised circular pieces of the genome after their removal from the linear counterpart. In some cases, we have clear evidences that these circular DNAs are maintained; such as double minutes in cancer cells and telometric circles.(Carroll et al.; Dilley et al.; Li and Miralles Fuste, 2017) More recently, a study demonstrated that deleted circular DNA elements can be transcribed into dsRNAs and processed further contributing to small-RNA-mediated genome reorganization (piRNAs) in *Paramecium*.(Allen et al.) Therefore, a study focused on eccDNA will facilitate a comprehensive understanding of DNA deletions and their potential biological consequences.

Recent studies focusing on eccDNA (Moller et al.; Shibata et al.) in mouse tissues and yeast have provided intriguing clues with respect to eccDNA sequence distributions, despite relying on methods that are susceptible to DNA sequence-or structure-dependent biases. Specifically, recently published bulk eccDNA isolation methods are prone to enrich for linear DNAs bearing repetitive sequences, e.g.(Wetmur and Davidson). Additionally, enrichment protocols in which DNA is amplified by rolling-circle techniques (although potentially useful in amplifying vanishingly small circular populations) have evident drawbacks with respect to several potential biases (size and topology).(Arakawa and Tomita; Fire and Xu) A rigorous approach to profile eccDNA should greatly advance the field towards comprehensive characterization of eccDNA loci (in normal and diseased tissues). To this end, we first maximized the robustness of circular DNA sequencing information to allow for extensive unbiased standardized profiling of eccDNAs. We then provided novel genome-wide “circulome” maps of a whole organism *(C. elegans)* and both healthy and diseased human tissues. We show that *(i)* genomic circular DNA repertoire is a function of cell type and state *(ii)* eccDNA-mediated deletions are present in both normal and diseased backgrounds *(iii)* a subset of eccDNAs map to several coding regions known to produce a diversity of protein isoforms.

## Materials and Methods

### C. elegans strains and maintenance

*C. elegans* were grown at 16 °C (unless specified) on agar plates with nematode growth medium (NGM) seeded with *Escherichia coli* strain OP50.(Brenner) Some strains were provided by the CGC, which is funded by NIH Office of Research Infrastructure Programs (P40 OD010440). Strains used are: wild type animals, VC2010 (PD1074), a clonal derivative of Brenner’s original *C. elegans* strain N2 (Brenner), *glp-1(e2141ts)*, (Austin and Kimble; Yochem and Greenwald) and fem-3(q20gf) (strain JK816).(Barton et al.)

### Spermatocytes isolation

Sperm were isolated from a synchronized population of *fem-3(q20gf)* at the permissible temperature according to (Gent et al.). Briefly, after multiple washes in M9 buffer (to remove bacterial contamination), the animals were diced with a razor blade in a glass dish under the microscope. The mixture of released spermatogenic cells and carcasses was filtered through a double layer of 10μm Nitex blotting cloth (Wildlife Supply) and washed three times in M9 before flash freezing in liquid Nitrogen. Some of the total spermatocytes samples were spun down for 20 minutes at (21,130 × g) and the pellet was discarded. The supernatant containing smaller spermatocytes were flash frozen in liquid N_2_.

### C. elegans Genomic DNA isolation

Sperm eccDNA was isolated from *fem-3(q20gf)* mutant strains (Barton et al.); this mutation converts a hermaphrodite to a sperm-only-producing strain. Predominantly somatic eccDNA was isolated from a *glp-1(e2141ts)* mutant strain (Austin and Kimble; Yochem and Greenwald); *glp-1* encodes a Notch signaling protein that induces the formation of a germ line in *C. elegans*. When shifted to a non-permissive temperature at the L1 larval stage, *glp-1(e2141ts)* animals produce animals with a full soma but with germline that is reduced by >99%.

To prepare eccDNA and control genomic DNA, pellets of whole animals (or sperm pellets) were incubated with occasional gentle mixing in “Worm lysis buffer” (0.1M Tris, 0.1M NaCl, 50mM EDTA, 1% SDS, pH=8.5) and 25 μg/mL Proteinase K (roche) for 1.5 hour at 62°C. Standard procedures were used for DNA isolation. Briefly, after an NaCl precipitation step, nucleic acids were ethanol-precipitated and treated with RNAse A (Roche) for 1 hour at 37 °C in TE (10mM Tris-HCl, 1mM EDTA, pH =8.0). After the addition of 1M ammonium acetate, DNA was purified by phenol-chloroform and ethanol precipitation steps. Genomic DNA pellets were resuspended in TE and stored at -20 °C.

### Fibroblast and granulocyte DNA isolation

For comparison of Fibroblast and Granulocyte eccDNA profiles, DNA from a previous whole genome sequencing study of a male individual (Merker et al.) was used. Fibroblast cells were derived from a punch biopsy of healthy skin, while granulocytes (present at high levels due to myelofibrosis) were isolated from blood. The subject (Merker et al.) was counseled and consented under a research protocol approved by the Stanford University Administrative Panel for the Protection of Human Subjects. (Merker et al.) DNA was extracted using the Gentra Puregene Cell Kit (Qiagen, Valencia, CA, USA) according to the manufacturer’s protocol.

### eccDNA enrichment

#### a. CsCl

High-molecular weight genomic DNA was mixed into 2.0 mL of a CsCl solution having a density of 1.55 g/mL. The sample was subjected to centrifugation at (500,237 × g) for 2.5 h in a S120-VT vertical rotor (Thermo Scientific). As a reference, a plasmid-DNA sample was run in parallel in a separate centrifuge tube. In the absence of exoV treatment, a distinct band, corresponding to sheared linear nuclear DNA (along with nicked/relaxed DNA circles), was visible under ultraviolet light. The plasmid-DNA control sample showed two distinct bands corresponding to nicked and linear (top) and covalently closed plasmid (bottom). A hypodermic needle was used to carefully isolate the fraction of interest with the closed circular plasmid band used as an indicator of the approximate location of the invisible eccDNA band. Ethidium was removed from the isolated bands by extraction with CsCl/TE-saturated 1-butanol. Samples were dialyzed for 2 days against TE buffer (10 mM Tris-Cl, 1mM Na_2_EDTA) at 4 °C.

#### b. ExoV

For enzymatic removal of linear DNA, 200 ng of Genomic DNA was treated with 400U/μg exoV (NEB) over 3 days in (50mM potassium acetate, 20mM Tris-acetate, 10mM magnesium acetate, 1mM DTT, pH=7.9) in the presence of 2 mM ATP, and 100μg/mL Ampicillin (to limit bacterial contamination). After each round of exoV treatment, the reaction was heat-inactivated at 70 °C for 30 min. A similar protocol was followed for human genomic DNA treatment, except for the duration of the reaction (5 days). For human eccDNA experiments, three different exoV to DNA ratios were used ranging from 500U/μg to 1000U/μg.

### Library prep and low-input nextera protocol

To generate fragmented genomic DNA libraries with appropriate linkers, 1 ng of DNA was treated with 1.5 ul of NexteraXT tagmentase (illumina) at 37 °C for 30 min with gentle shaking. For eccDNA libraries with DNA input lower than 1 ng, the eccDNA sample was treated with 0.5 ul of Nextera XT tagmentase at 37 °C for 30 min. In experiments where an enrichment for singly tagmented circles was sought, the tagmentation reaction was attenuated, with 0.5 ul of tagmentase incubated with the DNA at 37 °C (without shaking) for only 3 min. A minimal number of PCR cycles was chosen by monitoring the amplified DNA by gel electrophoresis after varying numbers of PCR rounds, ensuring libraries are prepared from PCR reactions in which the amplified DNA was still undergoing amplification. We find that amplification up to 10-12 cycles of PCR is sufficient for library production.

### Plasmids and synthetic DNA mini-circles

As a template for generating reference DNA circles, we used a plasmid with two directly repeated loxP sites cloned in the backbone of the generic vector pGEM5Zf(+).(Shoura and Levene; Shoura et al.) The resulting plasmid, pCS2DloxPzero(Shoura et al., 2012) allows insertion of arbitrary spacer sequences (gBlocks, IDT) between the two loxP sites by linearizing the plasmid with both NotI and PstI (NEB). Using pCS2DloxPZero as a vector, we cloned a 378 bp insert sequence between the two loxP sites, resulting in plasmid pCS2DloxP378. Upon treating pCS2DloxP378 with Cre recombinase (purified in house according to (Gelato et al.; Martin et al.)), a 412bp circle is produced (378 bp + one loxP site; 34 bp), along with the 3034 bp parent plasmid.

### Bioinformatic Analysis

Bowtie2 (version 2.2.25) was used align the paired end reads to the nematode (c10) or human (hg38) reference genomes respectively. Mapped reads were deduplicated using Picard. Unique reads were sorted and indexed using (samtools 1.2.). To analyze sequences that cannot be mapped uniquely, a separate positioning approach was used. This approach uses both unique and repeated k-mer sequences(Li et al.) to characterize individual k-mer/read counts and positions (python scripts available on request). Reads were divided into categories as follows: <u>Unique Chromosomal</u>: These reads represent the number of different read pair start/stop positional combinations for which both reads are uniquely and unambiguously mapped the reference genome (consistent with Bowtie algorithms) <u>Locally repeated reads or “focal repeats”</u>: defined as sequence that occur multiple times in the genome but for which all occurrences are confined to a single chromosome in a limited range of base-pair distance (chosen as 300kb for this study). <u>Dispersed Repeats</u>: Repeated reads that are distributed beyond this limited range or on multiple chromosomes. <u>Intrachromosomal Repeats:</u> defined as read that map to multiple sites on a single chromosome where the sites are separated by long distances (in this case, avove the arbitrary cutoff of 300Kb).

## Results and Discussion

### Circ-Seq as a hybrid biophysical-biochemical-bioinformatic method to characterize genomic circular DNAs

The approach described here entails two independent and effective separations of linear and circular DNA: *(i)* a subtractive biochemical, enrichment-based method: multiple rounds of extensive digestion with exonuclease V (removing the vast majority of linear DNA),(Palas and Kushner) and *(ii)* a biophysical purification method: centrifugation in CsCl/ethidium-bromide gradients, which provide topological separation of circular forms from linear DNA.(Grossman et al.) Applying one or both of these separations, we show a substantial enrichment for circular DNAs evidenced by enrichment for the circular mitochondrial DNA, which serves as an internal control, (See Figure 1). In both approaches, we avoid: (a.) DNA-purification methods that include denaturation and subsequent renaturation steps *(e.g*., based on alkaline lysis and neutralization), as these steps enrich for repetitive linear DNA fragments along with circles; (b.) digestion of genomic DNA with restriction enzymes or selection of a specific DNA size range through size-exclusion columns,(Shibata et al.) which naturally biases enrichment in favor of small eccDNAs; (c.) “rolling-circle” amplification of input DNAs to increase circular DNA copy number before subsequent processing. Notably, we find that that essentially identical populations of circular species are obtained using either method (i.) or method (ii.) (See Supplementary Figure 1 a and b)

**Figure 1.**
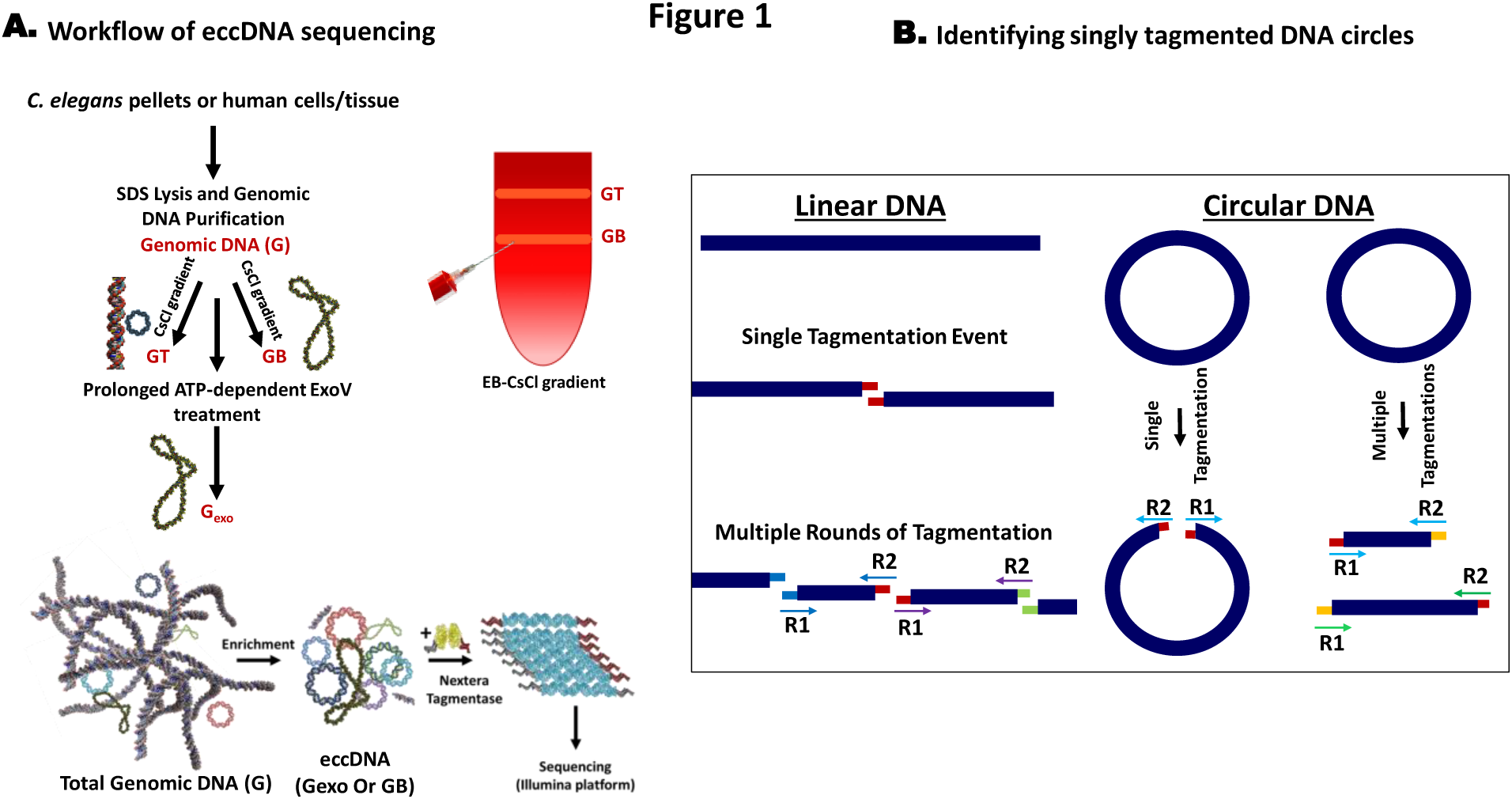
Workflow. **(A)** Genomic DNA is isolated from the organism/tissue of interest. Tissue is homogenized and treated with SDS and proteinase K. To enrich for circular DNAs, total genomic DNA (G) is treated with exoV (Palas and Kushner) to produce G_exo_ or banded in a CsCl gradient to separate G into GT and GB.(Grossman et al.) GT is the upper band of the gradient and includes linear DNAs and nicked circular DNAs. GB, the bottom band, consists of covalently closed-circular DNAs. After enrichment for circular DNA with either method (or both), eccDNA is minimally sheared by attenuated treatment with Nextera tagmentase. **(B)** Transposition creates a 9-bp sequence duplication flanking the transposon insertion site. Tn5 randomly binds and cuts DNA, leaving a staggered, 9-nucleotide single-stranded overhang. DNA on either side of the cut is filled by DNA polymerase in the first PCR cycle, thereby creating 9-bp duplications flanking the genomic DNA sequence. Matching overhangs in the figure have matching colors. Also, paired reads (R1 and R2, indicated by arrows) share the same color. If a circular DNA molecule gets cut only once by Tn5, paired-end sequencing will reveal a unique 9-bp duplication at the beginning of each read (designated by colored overhangs); thereby providing a bioinformatic mark for circular DNAs.

Following enrichment by procedure (i) or *(ii)* or both (i) and (ii), we find that eccDNAs can be simultaneously fragmented and tagged via a Tn5-transposition-based fragmentation and tagging system (Nextera tagmentase).(Caruccio; Reznikoff) The use of tagmentation, has the advantage of allowing us to work with very low levels of input material (< 1ng of eccDNA). An additional advantage of using Nextera fragmentation, in particular for small circles, is the duplication of a 9-nt segment of the target sequence on opposing sides of each transposon insertion.(Berg et al.) This feature provides a precise bioinformatic signature for the presence of singly tagmented circular DNAs in a sequenced eccDNA pool, (Figure 1b and Table 1) Our ability to capture circles in the protocol was confirmed with control circular substrates (3000-bp and 400-bp DNA circles) (Supplementary Figure 2). Extensive analysis of *C. elegans* eccDNA shows that eccDNA enrichment is captured in a quantitatively reproducible manner (Supplementary Figure 1). Confirming the specific role of exoV in eccDNA enrichment, we note that eccDNA is unenriched when ATP was omitted from the ATP-dependent exoV reactions, (Supplementary Figure 3).

**Figure 2.**
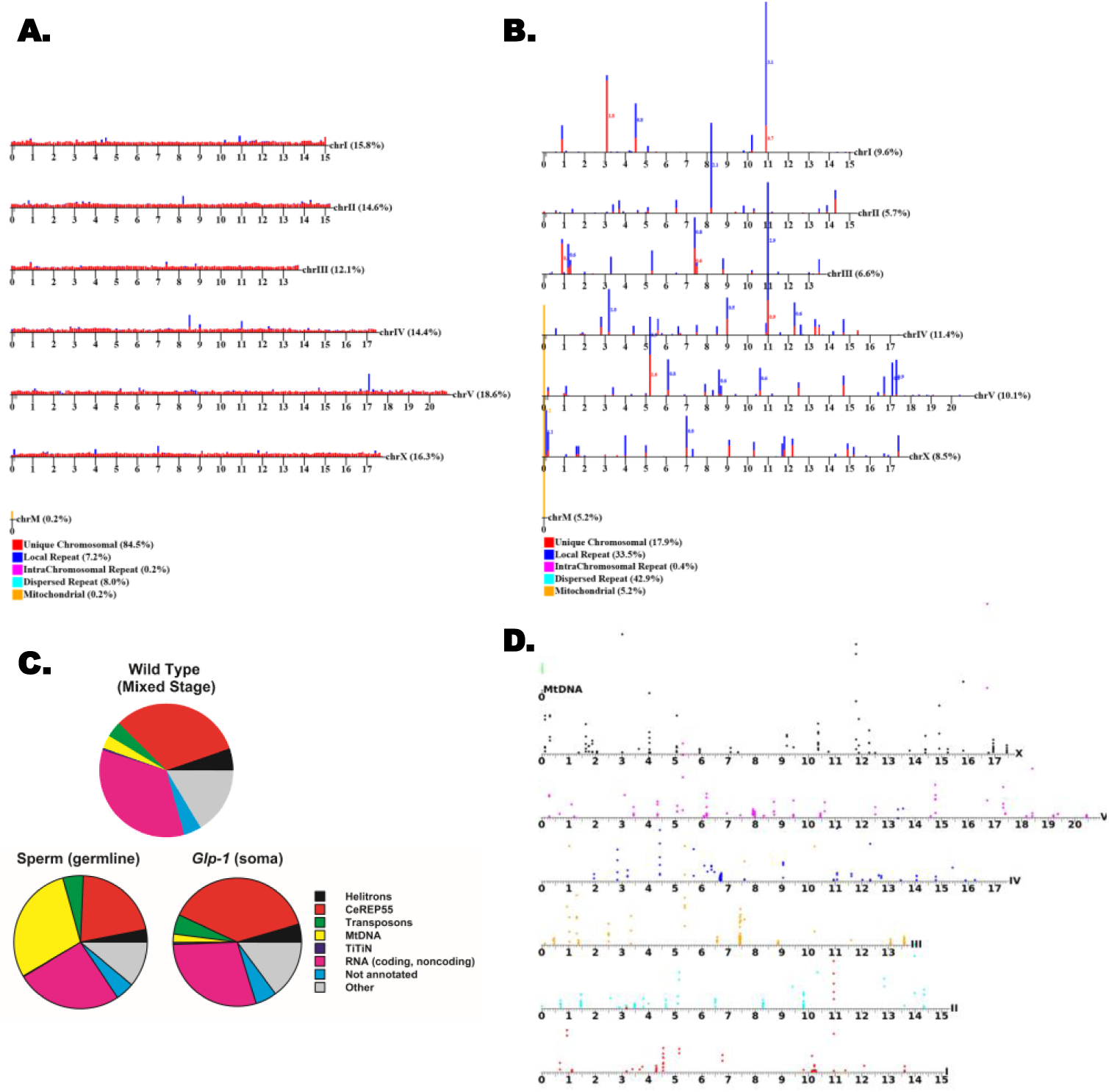
Data analysis. **(A)** and **(B)** are chromosomal maps of aligned reads in total genomic DNA (G) and eccDNA (G_exo_) respectively. Reads are categorized as: unique, local repeats, intrachromosomal repeats, and dispersed repeats. The graphs show unique reads and focal repeats only, as dispersed repeats cannot be mapped to one location. **(C)** Whole genome distribution of sequence classes in eccDNA fractions from WT animals, *C. elegans* sperm, and animals lacking germline cells **(D)** Our methodology applied to *glp-1* animals (somatic adults). This map shows uniquely mapped areas on each chromosome that are significantly enriched in the circular pool (1-kbp intervals with enrichment assessed through Bayes maximum-likelihood [minimum of 2-fold enrichment with a default false discovery rate of 0.05/(2*number of genes)]. This plot shows only reads that map uniquely to the genome. Position of the colored circle on the y-axis for each interval is proportional to the degree of enrichment.

**Supp. Figure 1.**
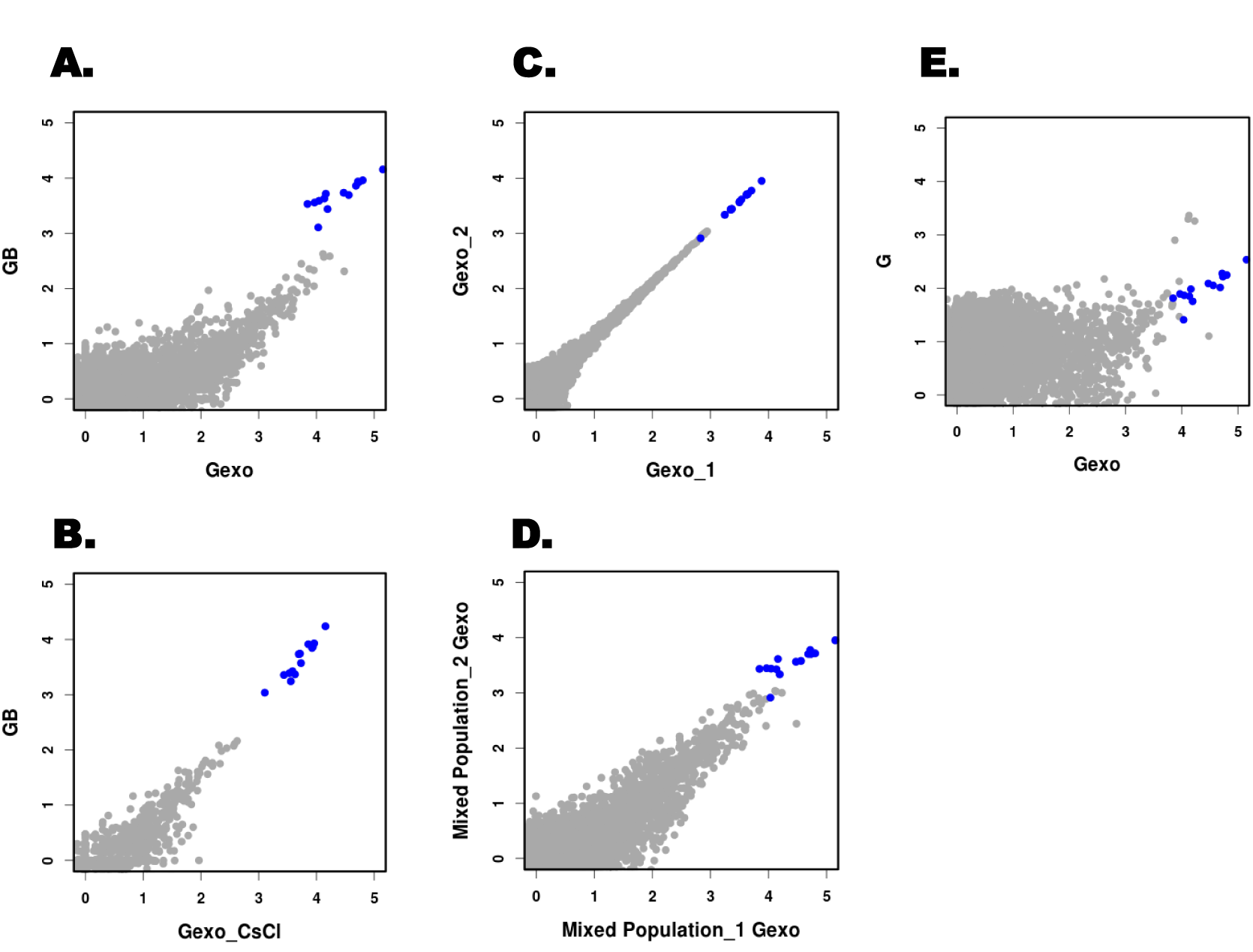
Reproducibility of eccDNA enrichment. **(A – E)** Plots showing log_10_ of read coverage for each chromosome with a bin size of 1000 bp. MtDNA is shown in blue. **(A)** compares eccDNA enrichment obtained by both methods (GB is the bottom band on a CsCl-EtdBr gradient and Gexo is total genomic DNA; G, treated with exoV). **(B)** compares GB to a Gexo sample purified on a CsCl-EtdBr gradient. **(C)** and **(D)** show data obtained for technical and biological replicates respectively. **(E)** shows the difference in coverage of total genomic DNA, G, in comparison to eccDNAs from the same sample, Gexo.

**Supp. Figure 2.**
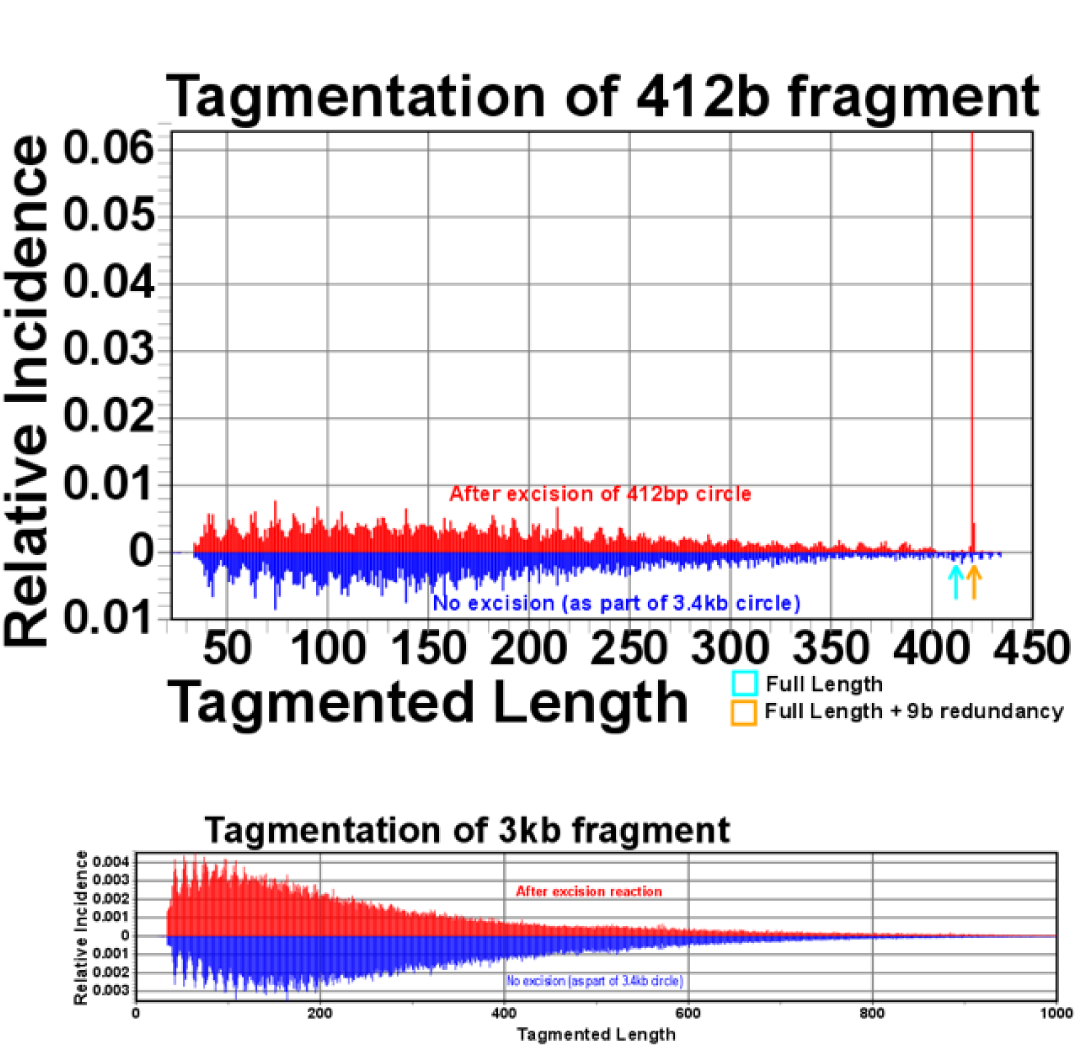
Testing tagmentation efficiency on circular DNA substrates of different length. A 3-kbp plasmid DNA (derived from pGEM5Zf+) and 412-bp circular DNA molecules were fragmented using Nextera-XT tagmentase. This graph provides a position-by-position summary of aligned reads. The orange arrow indicates a read length of a singly-tagmented circle, 421 bp (412 bp + 9 bp duplication)

**Supp. Figure 3.**
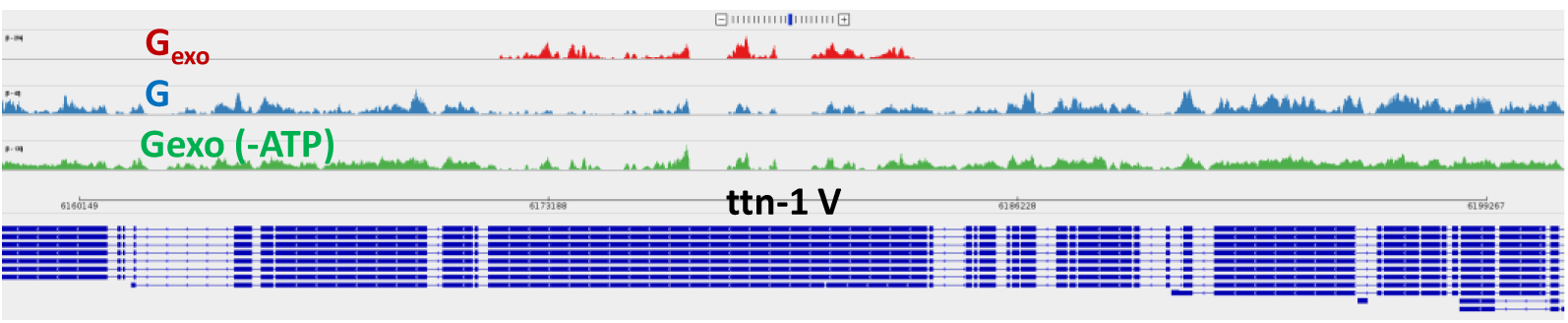
Enrichment of circular DNA is dependent on an optimal activity of exoV. Genomic DNA was treated with the same amount of exoV and for the same duration as described with the exception that ATP was not added to the reaction. The objective of this control experiment was to investigate biased enrichment of repetitive linear DNA due to PCR or other steps in the protocol. The observed enrichment of circular DNAs at the titin locus is lost when ATP was not added to the reaction *(i.e*., when exoV activity was prevented).

**Table 1.** Capturing singly-tagmented eccDNA circles

| Sample Name | Fraction of 9-bp duplication incidents relative to total captured incidents | Fold enrichment of singly tagged circles in eccDNA fractions over the corresponding total genomic DNA |
| --- | --- | --- |
| **Gexo** | 0.84 % | 311.1 |
| **GB** | 0.74 % | 277.8 |
| **Spermexo** | 0.22 % | 27.5 |
| **glp-1exo** | 0.4 % | 181.8 |
| **G** | 0.003 % | - |
| **SpermG** | 0.008 % | - |
| **glp-1G** | 0.002 % | - |

### eccDNA distribution in C. elegans somatic cells and germ line

To examine eccDNA distributions in a complex whole organism, we carried out circular DNA isolation and sequencing on eccDNA preparations from *C. elegans*. We investigated mixed stage *C. elegans* (whole worms), synchronized young larvae (L1 stage), synchronized germline-deficient adults *(glp-1* mutants, predominantly somatic tissue), and *C. elegans* sperm cells, (Supplementary Figure 4.) These analyses identified a diverse population of eccDNAs including segments from coding exons, transposons, repetitive regions, telomeric sequences, and other unannotated genomic locations (Figures 2 and 3). A substantial portion of eccDNA sequences originate from helitrons (a class of mobile elements known to transpose via a circular intermediate),(Kapitonov and Jurka) cut-and-paste transposons, and exons. (For a complete list of locations of eccDNAs and enrichment levels see Supplementary Data Set 1) Of these families, we were most surprised to observe circles directly derived from coding regions. Among the most abundant species are *ttn-1, plg-1, srap-1, clec-80, clec-223, frm-1, arrd-27, Y46B2A.3*, and *tag-80* genes *(ttn-1, Y46B2A.3*, and *tag-80* are the *C. elegans* orthologs of titin, mucin-1, and piccolo/aczonin respectively.) The *ttn-1* gene encodes a large protein that is essential in muscle function in *C. elegans*. Interestingly, the specific titin exon that is producing eccDNAs encodes an extended protein domain noted for its strong potential to form elastic structures of diverse lengths.(Guo et al.; Khan et al.; Werfel et al.)

**Figure 3.**
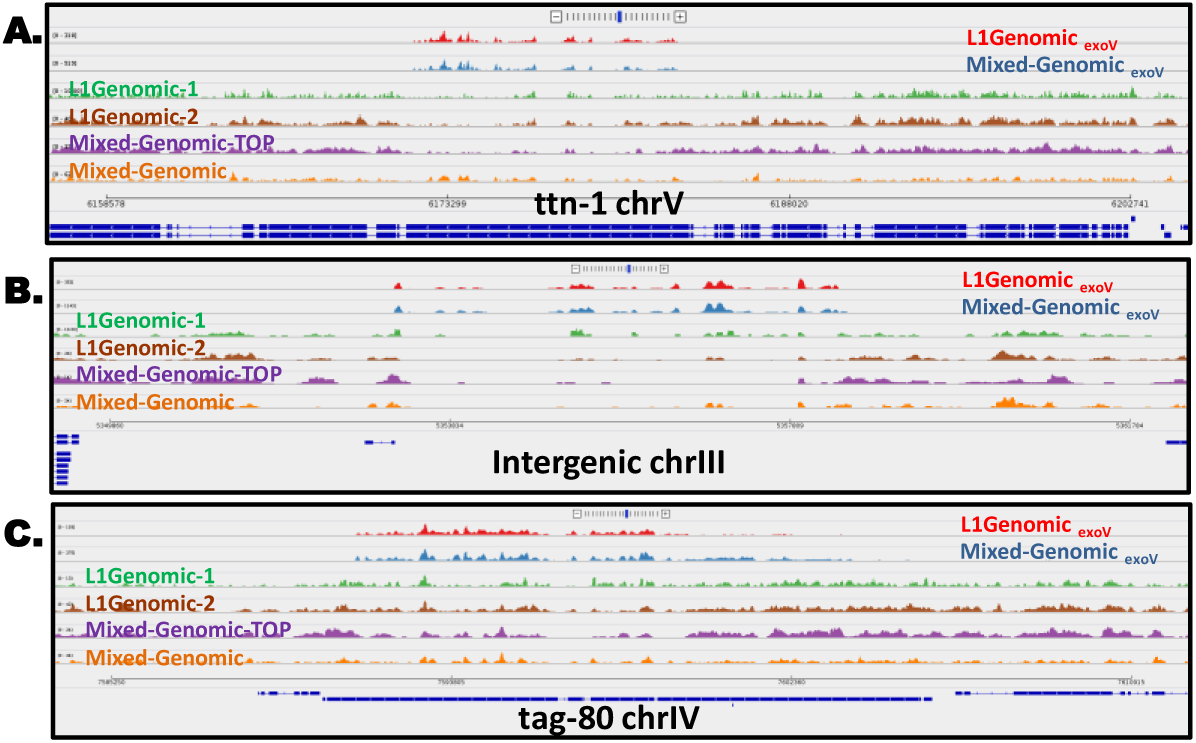
Sequence coverage of three eccDNA-enriched regions in different *C. elegans* populations. **(A) – (C)** show three distinct regions in the genome where eccDNA is generated. Red: exoV-treated DNA from synchronized young larvae, L1; blue: exoV-treated DNA from a mixed-stage population. Green, brown, and orange tracks are untreated genomic DNAs (L1 Genomic-1 and L1 Genomic-2 are independent biological replicas.). (A) Hyper-enriched eccDNAs isolated from exoV-treated *C. elegans* genomic DNA map precisely to a coding exon of the titin gene in the eccDNA pool. Unique mapping of untreated genomic DNA from the top band of a CsCl-EtdBr gradient, is shown in purple. **(B), (C)** show similar profiles for eccDNAs corresponding to an intergenic repeat and the *tag-80* gene, respectively.

**Supp. Figure 4.**
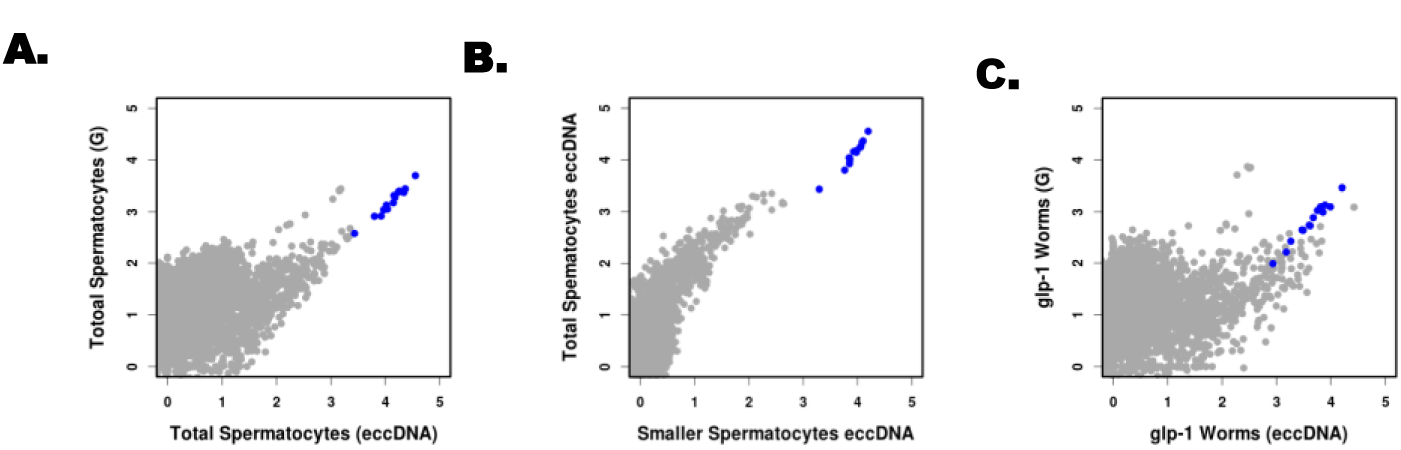
Analysis of eccDNAs from *glp-1* mutant animals and C. elegans spermatocytes. Plots showing log10 of read coverage for each chromosome with a bin size of 1000 bp. MtDNA is shown in blue. The top graph shows the difference in coverage of total genomic DNA from sperm versus sperm eccDNA. The bottom graph compared the coverage of total genomic DNA from *glp-1* animals versus eccDNA from the same sample. The middle graph compares eccDNA obtained from total sperm (total spermatocyte eccDNA) to the fraction of smaller spermatocytes separated by centrifugation (smaller spermatocyte eccDNA).

### Characterizing eccDNA distributions in three different human tissues

To characterize eccDNA populations in human cells we isolated eccDNA from human genomic DNA samples obtained from three sources: *(i)* a lymphoid cell line that has been subject to extensive sequence analysis and used as a standard for technical and software benchmarking in the genomics community (“Genome in a Bottle” cell line; GM12878),(Salit et al., 2014) *(ii)* neoplastic granulocytes from a patient with primary myelofibrosis, a subtype of myeloproliferative neoplasms (MPN), and *(iii)* a normal non-transformed primary fibroblast population from the same patient. This analysis showed extensive, but region-specific, eccDNA production (Figure 4). Overall, classes of sequences compared closely with those previously identified in the *C. elegans* genome such as coding and non-coding segments along with focal and dispersed repetitive sequences. Moreover, we find that a significant portion of the GM12878 “circul-ome” maps to mucin genes (such as *MUC1, MUC2, MUC6*, and *MUC17)* encoding high-molecular weight proteins characterized by the presence of large amino acid tandem repeat sequences that show allelic size variation.(Fowler et al.; Jia et al.; Linden et al.; Walsh et al.) A complete list of the eccDNA coordinates in GM12878 is presented in Supplementary Data Set 2.

**Figure 4.**
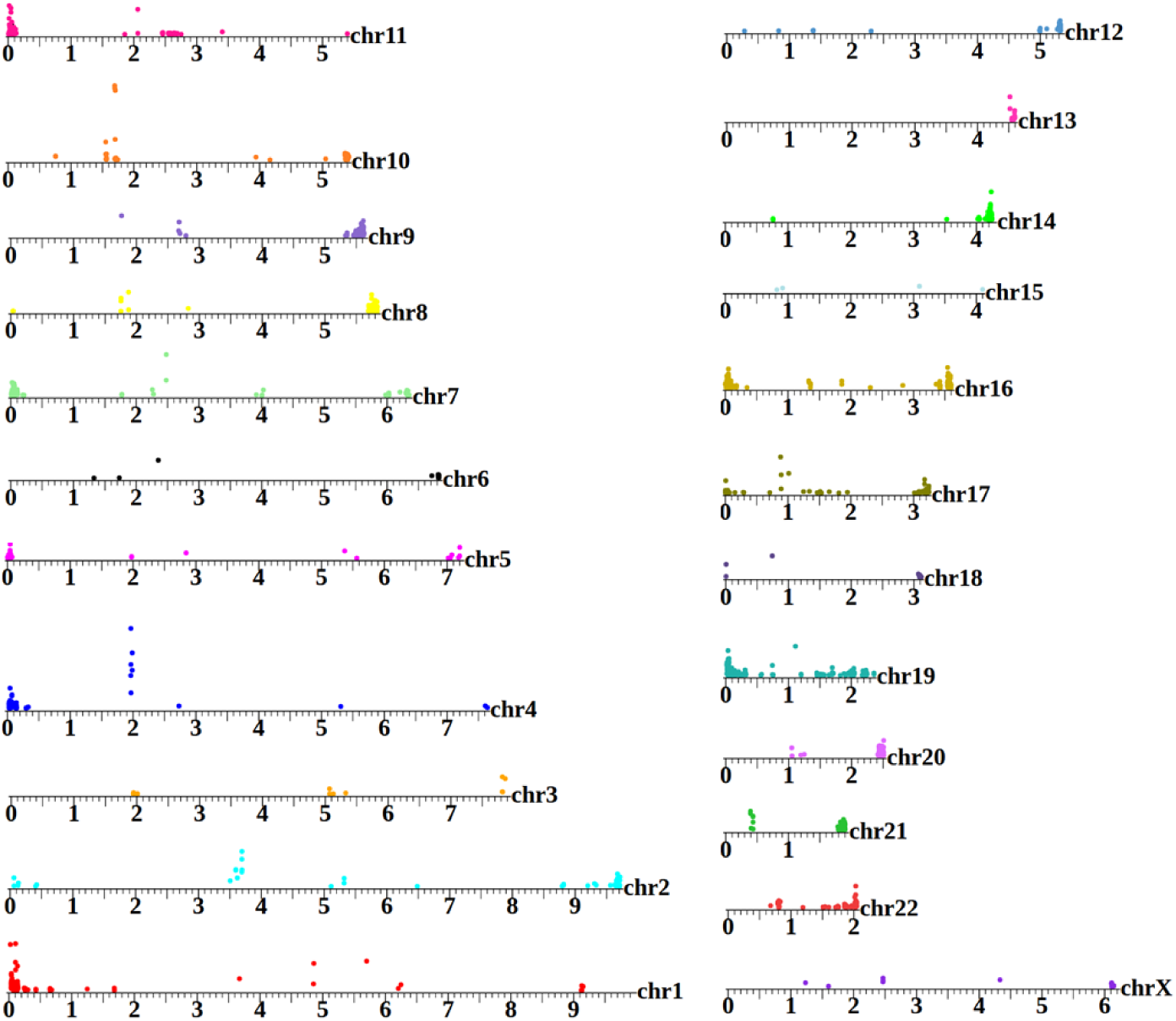
A human eccDNA map. GA12878 is the canonical cell line used by the National Institutes of Standards and Technology [NIST] as a benchmark for high throughput genome analysis.(Salit et al.) To date, published analysis of this sample to has, however, been focused on the linear genome. This map shows areas on each chromosome that are significantly enriched in the circular pool (25-kbp intervals with enrichment assessed through Bayes maximum-likelihood). Position of the colored circle on the y-axis for each interval is proportional to the degree of enrichment. The eccDNA profile of each chromosome is distinctive, with enriched regions aligning to coding segments, repetitive, and subtelomeric sequences. This plot shows only reads that map uniquely to the genome.

**Supp. Figure 5.**
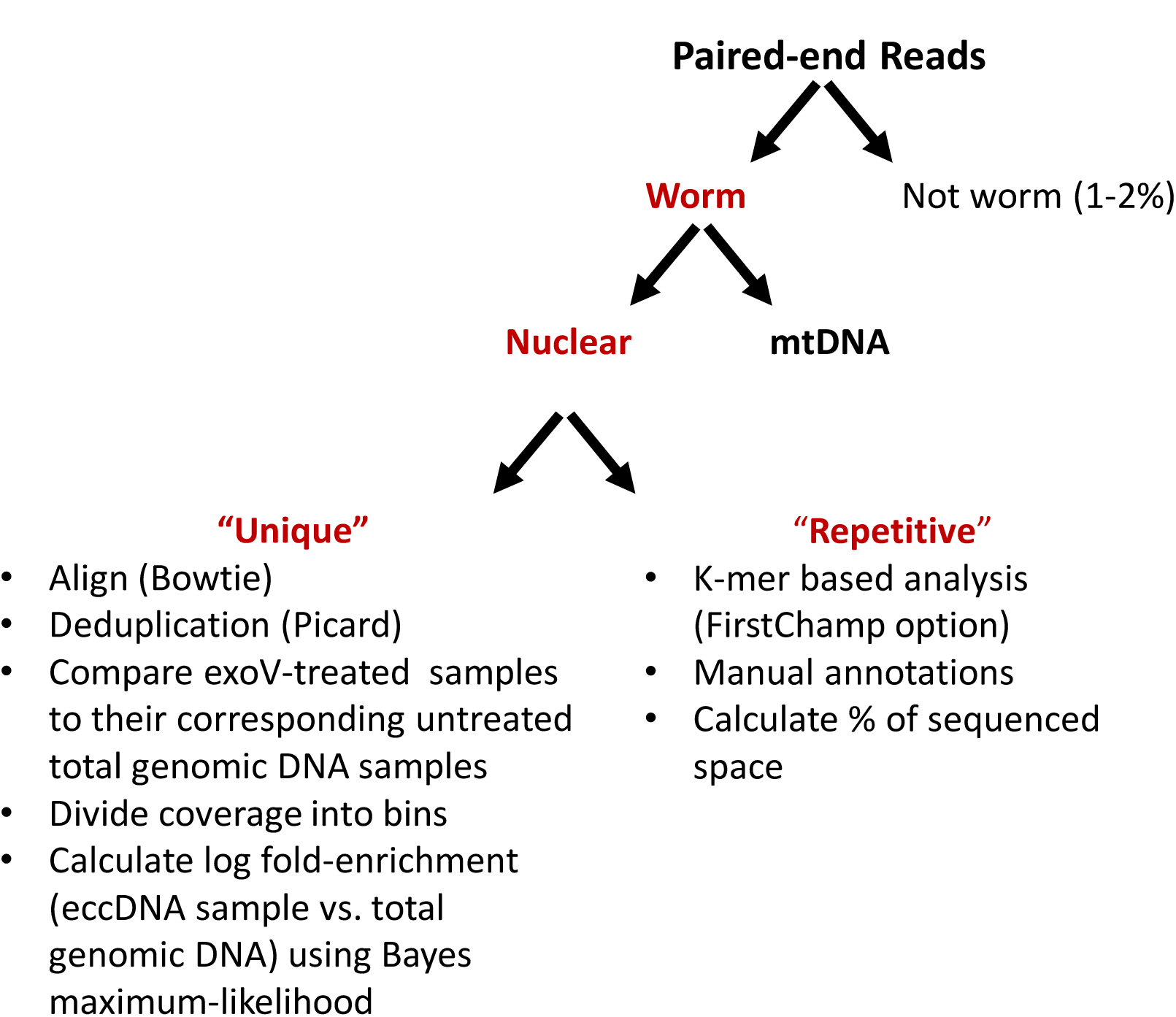
A schematic of data analysis. Paired-ends reads were analyzed using two independent pipelines described in the graph. k-mer analysis is used to analyze “repetitive reads” (reads that map to multiple locations in the genome). We chose to collapse all of the repatitve k-mers to the first alignable location in the genome (allowing for 2 mismatches); we named this approach “First Champion”. Unique reads were aligned with Bowtie in both eccDNA and total genomic DNA samples. Enrichment analysis was done through Bayes maximum-likelihood.

To evaluate the sensitivity and robustness of the human assay and to assess whether eccDNA profiles are cell specific, we compared circular DNA profiles between biological replicates of each cell type and between cell types. The eccDNA profiles obtained for each biological replicate pair are highly correlated, which indicates the reproducibility of the assay (Fig.5 A and B). Between the different cell types, we observe substantial differences, with cell state/type as a likely component in determining the diversity in the circul-ome. This holds true even when we compared eccDNAs in fibroblasts and granulocytes derived from the same donor (Figure 5).(Merker et al.)

**Figure 5.**
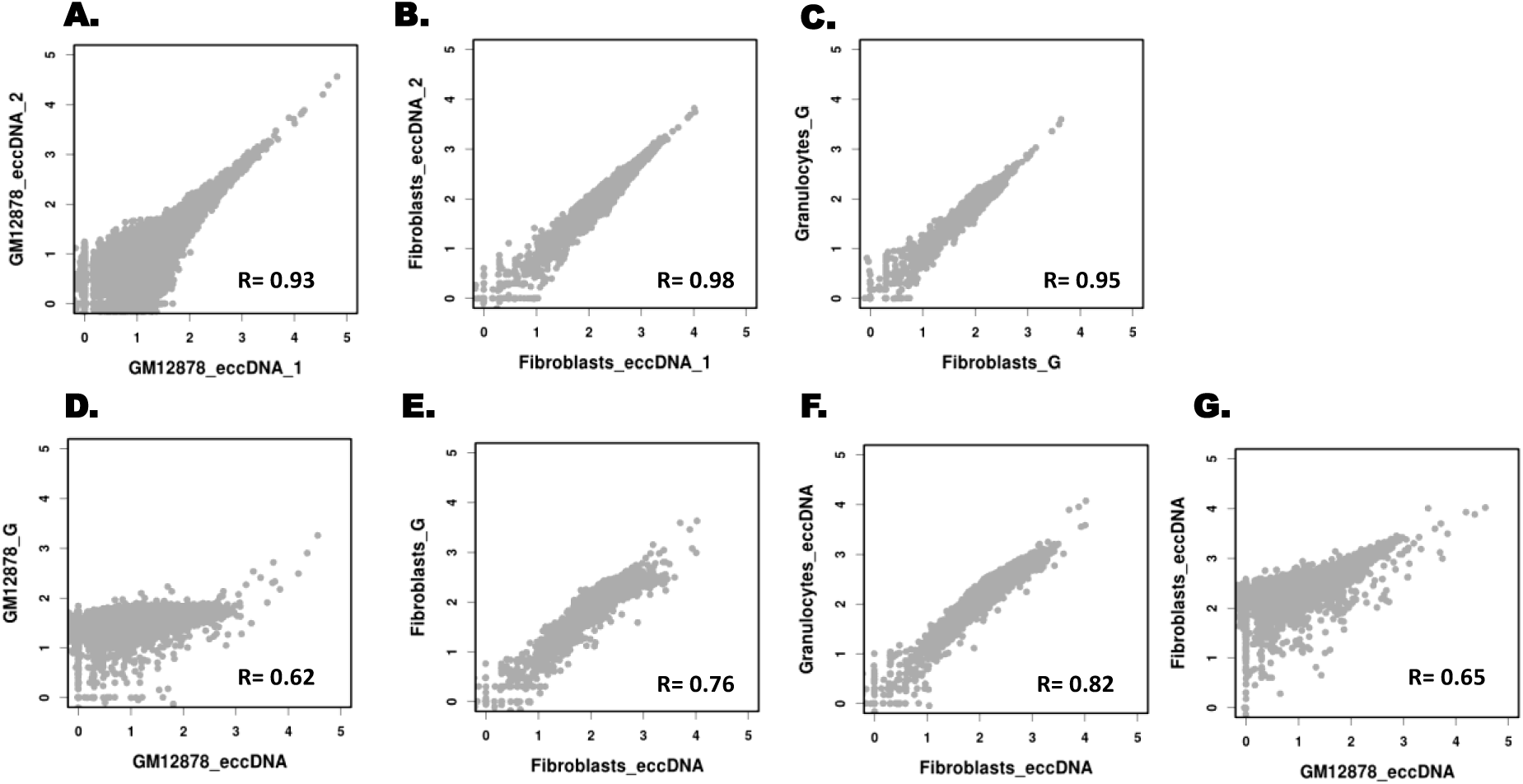
Analysis, reproducibility, and cell-type specificity of human eccDNAs. **(A - G)** showing log_10_ of read coverage for each chromosome with a bin size of 100 kbp. (A) and **(B)** show that eccDNA from the same tissue type is captured reproducibly. **(C)** Similar reproducibility is obtained for total genomic DNA (G) from different tissue types (normal fibroblasts *vs*. MPN granulocytes from the same individual). Distinct differences are evident when eccDNAs are compared to their reference total genomic DNAs **(D, E)** as well as when eccDNAs from different cell types are compared **(F, G)**.

In summary, we present a rigorous approach to isolating and purifying endogenous circular DNAs from *C. elegans* and human tissues. We have identified thousands of eccDNA regions in the genomes of *C. elegans* and human cells. Interestingly, the identified eccDNA species are enriched for specific exons that encode multi-isoform proteins (*e.g*., titin, mucins, and piccolo/aczonin.) A main finding of this study is that different cell types harbor different repertoires of circular DNAs. It has been shown that eccDNA copy number can be modulated by chromatin remodeling machinery.(Peng and Karpen) Therefore, we speculate that the circulome of a cell is a function of the genome’s unique and tandemly repeated sequence elements, recombination hotspots, and open chromatin.

## Acknowledgements

We thank the Fire laboratory for reading the manuscript; K. Artiles for technical support; and C. Smith and M. Bassik for valuable input. The use of high-performance computing resources of the FireLab Server and TARDIS is greatly acknowledged. This work is supported by grants from NIH (R01GM37706) (to AZF), Stanford Medicine Dean’s Postdoctoral Fellowship (to MJS), Human Frontiers Science Program Postdoctoral Fellowship LT000517/2011 (to IG), NIH/NSF Joint Program in Mathematical Biology (DMS-0800929) and Cecil and Ida Green Endowment (to SDL). The authors express gratitude to the Charles and Ann Johnson Foundation for their support of MPN research at Stanford. MJS and AZF conceived of the study. MS lead project development, designed experiments, developed the eccDNA assays, and performed all experiments. IG helped MJS set up *C. elegans* experiments in the initial phase of the project. MJS and AZF analyzed the data with input from LH and IG. JM and JG provided DNA from defined hematopeotic populations. SDL provided materials and experimental input. Overall discussions of the data and implications involved MS, IG, LH, SDL, and AZF. MJS, AZF, and SDL wrote the manuscript with input from all authors.

References

